# Multi-allele species reconstruction using ASTRAL

**DOI:** 10.1101/439489

**Authors:** Maryam Rabiee, Erfan Sayyari, Siavash Mirarab

## Abstract

Genome-wide phylogeny reconstruction is becoming increasingly common, and one driving factor behind these phylogenomic studies is the promise that the potential discordance between gene trees and the species tree can be modeled. Incomplete lineage sorting is one cause of discordance that bridges population genetic and phylogenetic processes. ASTRAL is a species tree reconstruction method that seeks to find the tree with minimum quartet distance to an input set of inferred gene trees. However, the published ASTRAL algorithm only works with one sample per species. To account for polymorphisms in present-day species, one can sample multiple individuals per species to create multi-allele datasets. Here, we introduce how ASTRAL can handle multi-allele datasets. We show that the quartet-based optimization problem extends naturally, and we introduce heuristic methods for building the search space specifically for the case of multi-individual datasets. We study the accuracy and scalability of the multi-individual version of ASTRAL-III using extensive simulation studies and compare it to NJst, the only other scalable method that can handle these datasets. We do not find strong evidence that using multiple individuals dramatically improves accuracy. When we study the trade-off between sampling more genes versus more individuals, we find that sampling more genes is more effective than sampling more individuals, even under conditions that we study where trees are shallow (median length: ≈ 1*N_e_*) and ILS is extremely high.

## 1. Introduction

Using a large number of loci to reconstruct the species phylogeny is becoming routine practice [1, 2, 3]. Beyond dramatically increasing the amount of data available [4], whole-genomes have enabled us to study individual genealogies, which can be discordant with each other and with the species tree [5]. Incomplete lineage sorting (ILS) is a major cause of such discordance [5, 6], and the multi-species coalescent (MSC) model [7, 8] has enabled a probabilistic study of ILS. In the presence of ILS, the traditional approach of concatenating data from multiple genes can become misleading [9, 10, 11], which has motivated researchers to develop alternative methods [12, 10]. One approach is to use a full Bayesian analysis under the MSC model [13, 14] to co-estimate gene trees and the species tree, but such analyses have severe computational limitations [15, 16, 17], and improving their scalability is a subject of active research [18, 19]. A more scalable approach first estimates gene trees individually, and then summarizes them to obtain the species tree, while accounting for the distribution on gene trees defined under the MSC model [20]; many such summary methods have been developed, including STAR/STEAC [21], NJst [22], STEM [23], GLASS [24], MP-EST [25], and more recently, ASTRAL [26]. Yet a third category of methods avoid gene trees altogether and estimate species trees from concatenated sequence data directly while accounting for gene tree discordance. Examples of this category of methods includes SNAPP [27], SVDQuartets [28], the PoMo model [29] and its implementation in IQ-Tree [30].

ILS can arise when multiple alleles of a gene survive through consecutive speciation events. For example, imagine an ancestral species (or population) *R* that gives birth to three present day species *X*, *Y*, and *Z*, with *X* and *Y* sharing a common ancestor *P*. Consider a locus with two alleles *a* and *b* in *R*. By random chance, only *b* survives in *Z*, whereas both alleles remain present throughout the life of *P*; however, only *a* survives in *X* and only *b* survives in *Y*. Such a scenario will create a linage tree (which we call a gene tree) that puts *Z* and *Y* as sisters to the exclusion of *X*, and this will be in conflict with the species tree that puts *X* and *Y* together. The probability of such a scenario will be much higher if *P* spans a relatively small number of generations or if it has a high effective population size [7]; the ratio of the number of generations to the haploid effective population size is called the coalescent unit (CU) and gives a measure of branch length that directly relates to the expected amount of ILS [31]. A similar scenario is that *a* and *b* both survive in all five populations, but the single individual chosen to represent *X* happens to be homozygous for *a*, whereas individuals representing *Y* and *Z* happen to be both homozygous for *b*; this will also lead to a lineage tree that is discordant with the species tree. The likelihood of this scenario, in addition to being affected by the internal branch lengths, also depends on terminal branches. This second scenario of discordance makes it clear that for short terminal branches, the choice of individuals representing a species may matter.

To account for potential impacts of polymorphism in present-day species, several authors have suggested sampling multiple individuals per species (and/or phasing) to create multi-allele datasets where each species can have multiple alleles per locus [12, 14, 15, 32, 33, 34]. Models of sequence evolution that directly account for polymorphisms have been also developed [29]. Alternatively, one can use summary methods to analyze multi-allele datasets. This requires that gene trees estimated from sequence data are either multi-labeled by species names (i.e., several nodes in a gene tree are labeled by the same species) or are labeled by the name of individuals and a mapping between individuals and species is known. Then, the summary method can estimate the best species tree labeled with names of species; this is equivalent to finding the best species tree labeled with individual names constrained to each species being monophyletic. Predefining species boundaries side-steps difficulties of defining boundaries of recently diverged species [35, 36], leaving that question to the analyst. In other words, the approach assumes the species are correctly delimited.

When species boundaries are known *a priori*, the MSC model easily extends to the multi-individual case [37, 38, 39]. While the evidence for the cost-effectiveness of sampling multiple individuals remains mixed [40], several methods exist that can use such data [14, 22, 25]. To our knowledge, NJst is the only summary method that can handle multi-individual *unrooted* gene trees [22], and after a fix in handling multi-individual data [41], the NJst method is now statistically consistent for multi-individual datasets.

One commonly used method of species tree reconstruction is ASTRAL [26, 42], which has been used on many phylogenomic datasets. Given a set of unrooted input gene trees, ASTRAL seeks to compute the species tree that shares the maximum total number of induced quartets with the input set of gene trees. A constrained version of this NP-hard problem [43] is solved by ASTRAL in polynomial time, but solutions to the constrained problem are proved statistically consistent under the MSC model [26].

The published ASTRAL algorithm, until now, could only take single-labeled trees as input and had no way of handling multi-allele datasets. Incipient implementations of a feature in ASTRAL to handle multi-allele datasets were not rigorously tested; nor were they formally described. In this paper, we will introduce a new algorithm for handling multi-labeled gene trees in ASTRAL (which is different from the previous untested method) and establish its accuracy on both simulated and empirical datasets. We show that the quartet optimization problem extends in a natural way to multi-labeled datasets, leaving us with only one difficulty: defining the constrained search space for ASTRAL. We propose and test heuristic approaches based on subsampling individuals to build a sufficiently large search space. We test the method in extensive simulations and ask whether using multiple individuals results in improved accuracy compared to having single individuals. We also test if predefining species boundaries improves accuracy and compare the accuracy of ASTRAL to NJst.

## 2. Theory

### 2.1. Background on ASTRAL

We are given a set *𝒢* of *k* unrooted gene trees, singly-labeled by the leaf-set *L* of *n* taxa. *There are* 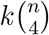 quartet trees induced by the input set *𝒢*. The Weighted Quartet (WQ) score of any candidate species tree is defined as the number of the 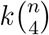 quartet trees that the candidate tree also induces. ASTRAL seeks to find the species tree that maximizes the WQ score [26, 42].

At the heart of the ASTRAL algorithm is the ability to score a tripartition of the species leaf-set in isolation from others tripartitions, which then enables the use of dynamic programming. The dynamic programming starts from the set *L* and recursively divides it into smaller subsets, each time choosing a division that maximizes the number of shared quartets. If we consider all ways of dividing a subset into smaller subsets, the problem is solved exactly but in exponential time. To obtain a polynomial time algorithm, we define a constraint set *X* of bipartitions and restrict the search to tripartitions derived from the set *X*. Let *X′* = {*A*: *A|L − A* ∈ *X*}. To constrain the search, we only consider divisions of a subset into two parts such that both parts appear in *X′*. We define *V*(*A*) as the score for an optimal subtree on *A*, and set *V*(*A*) = 0 for *|A|* = 1. Then, the dynamic programming recursion is:

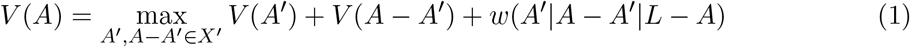

where *w* computes the number of gene tree quartet topologies that match any species tree that includes the tripartition *A′*|*A* − *A′*|*L* − *A* (see ASTRAL-II [42] – Eq. 2). ASTRAL-II, starts by including in *X* the set of bipartitions observed in the input gene trees and then supplements that set using a set of various heuristics [42]; ASTRAL-III slightly changes those heuristics such that the size of the set *X* is guaranteed to grow no more than linearly with both *n* and *k* [44].

### 2.2. Quartet score for multiple individuals

In the presence of multiple individuals, the definition of the quartet score can be easily generalized. Let *S* = {1… *n*} be the set of species and let *R* = {1… *m*} be the set of individuals. The input is a mapping *m*: *R → S* from individuals to species and a set {*t*_1_… *t_k_*} of unrooted gene trees, each labeled by *R_i_* ⊂ *R*. Following Allman *et al.* [39], for any species tree *T* labeled by *S*, we define an extended species tree *T_ext_*, labeled by *R*. The extended tree is built by adding to each leaf of *T* all individuals corresponding to that species as a polytomy (Fig. 1); i.e., for leaf *s* ∈ *S* of *T*, add a child *r* ∈ *R* for every *r* ∈ *m^−^*^1^(*s*)). We define the quartet score of an unrooted species tree *T* labeled by *S* with respect to the input gene trees as the quartet score of the extended species tree with respect to gene tree:

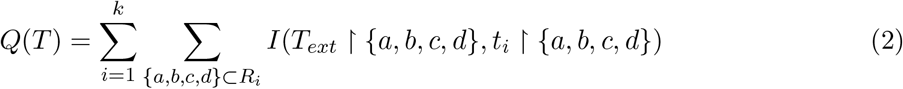

where *I*(*t*_1_*, t*_2_) indicates if its arguments are topologically identical trees and *t* ↾ *A* denotes the tree *t* restricted to the set *A*. ASTRAL in multi-individual mode seeks to find the species tree that maximizes this quartet score.

**Figure 1:**
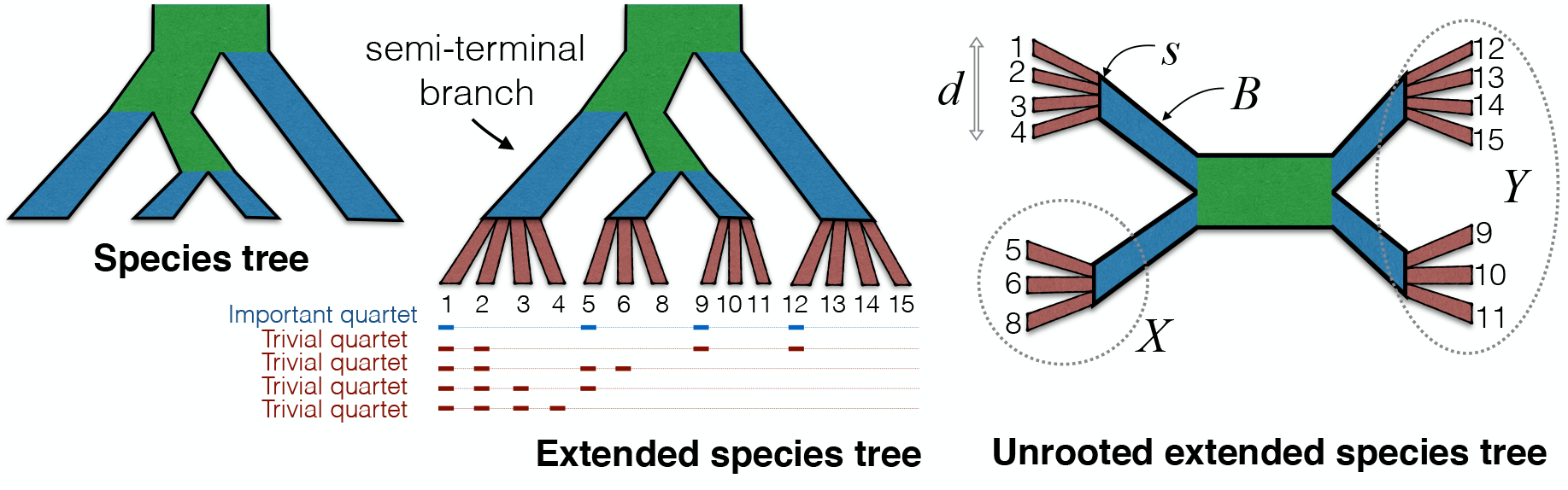
The illustration of rooted and unrooted extended species tree. Left: the species tree. Middle: the extended species tree with polytomies added for individuals of each species. Only quartets that have individuals from four species are important; the remaining quartets are trivial. For the top two trivial quartets, the extended species tree induces a resolved tree but all valid extended species trees define *the same* tree and therefore the quartet does not help in deciding among species trees. For the bottom two trivial quartets, all extended species trees give an unresolved quartet, and therefore, do not help in distinguishing species trees. Right: the unrooted extended species tree. The species *s* has *d* individuals; *B* is a semi-terminal branch in the extended species tree and corresponds to the terminal branch of *s* in the original species tree. Removing *B* and its two adjacent branches divides the tree into three groups, one corresponding to *s* and two opposite groups shown here as *X* and *Y*. Green: internal branches, Brown: branches corresponding to individuals, Blue: terminal/semi-terminal branches.

To show the connection between the quartet score and ILS, we show that ASTRAL is a statistically consistent estimator of the species tree for multi-individual randomly-sampled error-free gene trees given a correct mapping between species and individuals. We do so using results by Allman *et al.* [39] who showed (Corollary 10 [39]) that a coalescent process on the extended tree with one sample per species leads to exactly the same distribution of gene trees as the the multiple-individual process on the original species tree.

The set of all (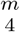) quartets can be divided into two types: trivial and important quartets. A quartet *Q* = {*a, b, c, d*} is trivial if any gene tree reduced to *Q* either contradicts every extended species tree or contradicts no extended species tree; in contrast, a gene tree resolution of an important quartet matches some extended species trees and contradicts others. Let *m*\*(*X*) = {*m*(*r*)|*r* ∈ *X*} give the set of species for a set of individuals. It’s easy to see that a quartet *Q* is important if |*m*\*(*Q*)| = 4; i.e., if it includes at most a single individual from any species. For example, in Figure 1, the quartet {1, 5, 9, 12} is important. All other quartets (i.e., those that include at least one species with multiple individuals) are trivial. To see this, first consider |*m*\*(*Q*)| = 3: two individuals (say, *a* and *b*) chosen from the same species and the other two chosen from two different species (e.g., {1, 2, 9, 12} in Fig 1). Any unrooted gene tree that puts *a* and *b* on one side of a quartet and *c* and *d* on the other side will match every possible extended species tree. Similarly, every gene tree that puts *a* and *b* on the opposite side of a quartet tree will contradict any possible extended species tree. Thus, these quartets are trivial. A similar argument carries if two individual are from one species and the other from a second species (|*m*\*(*Q*)| = 2; e.g., {1, 2, 5, 6} in Fig 1)). Quartets that include three (e.g., {1, 2, 3, 5} in Fig 1) or four (e.g., {1, 2, 3, 4} in Fig 1) individuals from the same species will always be unresolved in any extended species tree and thus cannot contradict any extended species tree. Thus, these are also trivial.

For all important quartets, since one individual is chosen per species, the unrooted species tree topology matches the most probable unrooted gene tree [39]. Thus, trivial quartets are inconsequential in the choice of the species tree and for important quartets, the central results required for proving ASTRAL statistically consistent carries through to multi-individual datasets. Therefore, the same argument used to prove ASTRAL statistically consistent for the single individual datasets [26] can be used here to argue that ASTRAL is statistically consistent.

### 2.3. Dynamic programming algorithm for multi-individual datasets

To find the species tree that maximizes the quartet score, we can continue to use the dynamic programming given in Equation (1) with an additional constraint: individuals of a species should not be separated into two parts. Thus, the new dynamic programming recursion is still given by equation (1) with the two changes: i) the set *X′* never includes a cluster (half a bipartition) that has some but not all individuals of a species; (ii) the boundary condition is changed to |*m*\*(*A*)| = 1. In other words, we stop as soon as *A* includes only individuals from a single species. With these constraints, for any *w*(*X*|*Y*|*Z*) calculation, all individuals of any species will belong to only one of *X*, *Y*, or *Z*. The dynamic programming produces the extended species tree, which can be simply mapped to a *S*-labeled species tree by removing all terminal branches.

Because the dynamic programming stops short of resolving the relationship between individuals of the same species, some quartets will not be counted by the dynamic programming. Note that to find the species tree that maximizes the score, it does not matter if we count a given trivial quartet, as long as we either always count it or always ignore it. Important quartets, however, should always be counted. A gene tree quartet topology *ab/cd* is counted by the *w*(*X*|*Y*|*Z*) function if there exists a permutation of (*X, Y, Z*) denoted as (*U*_1_*, U*_2_, *U*_3_) so that *a* ∈ *U*_1_, *b* ∈ *U*_2_ and *c, d* ∈ *U*_3_ or *c* ∈ *U*_1_, *d* ∈ *U*_2_ and *a, b* ∈ *U*_3_. It’s easy to see that an important quartet will be counted by the dynamic programming exactly twice, just like single individual ASTRAL, because the constraints have no bearing on quartets with one individual from four different species.

The case of trivial quartets is more complicated. A quartet *Q* with individuals from two or one species (|*m\**(*Q*)| ≤ 2) will always intersect with at most two of sides of *X*|*Y* |*Z*, and thus, can never be counted by any *W* (*X*|*Y*|*Z*) calculation. This leaves us with one form of trivial quartets, namely, those that includes individuals from three species. Let *Q* = {*a*_1_*, a*_2_*, b, c*} where *m*(*a*_1_) = *m*(*a*_2_) ≠ *m*(*b*_1_) ≠ *m*(*c*_1_). For any species considered by the search space of the dynamic programming, there exist a tripartition *A′*|*A* − *A′*|*L* − *A* scored in Equation 1 where *a*_1_ and *a*_2_ belong to one side, *b* belongs to a different side, and *c* belongs to the third side and thus *w*(*A′*|*A* − *A′*|*L* − *A*) will count the number of genes where *Q* is resolved as *a*_1_*a*_2_/*bc*. Note that this is true for any tree that the dynamic programming could produce. Moreover, these quartets will be counted only once since there could never be a tripartition where *b* and *c* belong to one side, *a*_1_ to another side, and *a*_2_ to a third side. Thus these trivial quartets will contribute to the score of every species tree equally and will thus be inconsequential in the choice of the species tree.

To summarize, we showed that the dynamic programming, with two simple modifications will optimize the quartet score correctly. However, it will not compute the correct quartet score because it fails to count some trivial quartets. To address this, at the end, the score of the species tree is recomputed with a simple procedure that explicitly counts trivial quartets.

### 2.4. Defining the search space

The final and the main difficulty in using multi-labeled gene trees is defining the search space, which entails constructing the constrained set *X*. Recall that any *A* ∈ *X* should include either all or none of the individuals of any species. Simply adding bipartitions from the gene trees, as ASTRAL-I, ASTRAL-II, and ASTRAL-III all do, will violate this constraint. In order to address this, for each gene tree *t_i_*, we create 100 singly-labeled gene trees 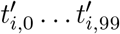 by randomly choosing one individual from each species, taking the induced tree, and renaming leaves from *R* to *S* using the mapping *m*. Then, we compute the greedy consensus of 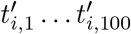, which will be labeled by *S*, and we extend it (just like building an extended species tree) to get the tree 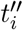 on leaf-set *R*. We proceed to build the set *X* using the usual ASTRAL-III approach with 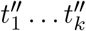 as input. Thus, the set *X* will includes all bipartitions of 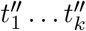 as well as selected resolutions of their polytomies; moreover, ASTRAL-III heuristics expand the set *X* using several methods [42, 44], many of which rely on a similarity matrix. To build the similarity matrix for pairs of species, we first build the ASTRAL similarity matrix [42] for pairs of individuals and then compute averages for all pairs of individuals corresponding to each pair of species as shown in Algorithm S1. To have a sufficiently large search space, we repeat the entire process for *r* rounds, setting *r* by default to 1 + ⌈log_2_ max_*i*∈*S*_ |{*j* ∈ *R*: *m*(*j*) = *i*}|⌉. This default setting, which the user can adjust if desired, starts with 1 rounds if all species have a single species and slowly (i.e., logarithmically) increases the number of rounds as more individuals become available. For example, if the maximum number of individuals per species is 5, 10, 20, or 40, the number of rounds is set to 4, 5, 6, or 7, respectively. This slow pace of growth is based on our tests that shows the set *X* is sufficiently large with these settings.

### 2.5. Branch length and branch local posterior probability

One benefit of having multi-individual datasets is that the branch length for terminal branches can also be computed because terminal branches are internal branches in the extended species tree. We extend the approach currently used in ASTRAL [45], which estimate the ML branch length as 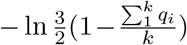; here, *q_i_* is the fraction of quartets defined “around” the branch of interest that agree with the species tree topology in gene tree *t_i_*. We have defined a quartet to be around an internal branch *b* if each of its four leaves come from the subtree attached to one of the four branches adjacent to *b*. For multi-individual datasets, let a species *s* have *d* = *|m^−^*^1^(*s*)| individuals. For the terminal branch *B* corresponding to the species *s* in the unrooted extended species tree, let *X* and *Y* be the two sides of the opposite sides of the branch *B* (see Fig. 1). There are 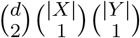 quartets such that the length of the quartet tree correspond exactly to the length of *B* (e.g., {1, 2, 5, 9} in Fig 1). We define *q_i_* as the proportion of these quartets that agree with the species tree in gene tree *i*. A naïve enumeration of all possible quartets would result in slowed computation of branch lengths; however, an algorithmic trick can reduce the running time. For each individual *i*, we create a quadripartition of leaves: {*i*}|*m*^−1^(*s*) − {*i*}|*X*|*Y*. For each quadripartition, we compute the branch length using the same fast technique introduced in a previous paper [45] for internal branches. This will require only *d* calculations per terminal branch; at the end, we have counted each quartet exactly *d* times, so we divide the sum by *d*. Also note that given *q_i_*, not only we can compute the branch length, but we can also compute the local posterior probability (localPP) of the terminal branches [45]. This measure can be interpreted as the probability of the predefined boundaries between species being in fact correct.

## 3. Materials and Methods

### 3.1. Dataset

We simulate two new datasets (see Table 1): a heterogeneous dataset (D1) where many parameters are simultaneously changed and ILS levels are extremely high and a more homogeneous dataset (D2) where parameters are less varied and the amount of ILS is controlled to create three model conditions. For both datasets, we use SimPhy [46] to generate species trees according to the birth/death model and gene trees according to the MSC model (exact parameters described in Appendix B). Each replicate of the simulation has its own species tree and all replicates have 5 individuals per species.

**Table 1:** Summary of properties of datasets D1 and D2. For parameters drawn from distributions other than uniform and for quartet scores and gene tree error, we show summary statistics: (5% percentile, mean, 95% percentile). For sequence length, distribution parameters are drawn from another distribution as detailed in Appendix B. *M* indicates 10^6^.

| Dataset | D1 (heterogeneous) | D2 (homogeneous) |
| --- | --- | --- |
| Number of replicates | 330 | 150 |
| Number of species | uniform(20,200) | 200 |
| Number of genes | uniform(50,1000) | 1000 |
| Population size | uniform($10^5$, $10^6$) | $2 \times 10^5$ |
| Birth-rate | log-uniform($10^{-7}$, $10^{-6}$)<br>($1.1 \times 10^{-7}$, $3.8 \times 10^{-7}$, $8.3 \times 10^{-7}$) | $10^{-6}$ |
| Death-rate | log-uniform($10^{-7}$, birth-rate)<br>($1.0 \times 10^{-7}$, $2.0 \times 10^{-7}$, $4.4 \times 10^{-7}$) | 0 |
| Number of generations | log-normal(13,0.5)<br>($1.9 \times 10^5$, $5.0 \times 10^5$, $1.0 \times 10^5$) | 0.5M, 1M, or 2M<br>(50 replicates each) |
| Sequence length | log-normal(-)<br>(406bp, 1096bp, 1803bp) | log-normal(-)<br>(322bp, 721bp, 1217bp) |
| Quartet score (ILS) | (0.37, 0.50, 0.75) | 0.5M: (0.48, 0.50, 0.53)<br>1M: (0.56, 0.62, 0.66)<br>2M: (0.72, 0.78, 0.83) |
| Average gene tree error (AGTE);<br>RF distance | (0.19, 0.4, 0.68) | 0.5M: (0.13, 0.42, 0.84)<br>1M: (0.10, 0.31, 0.77)<br>2M: (0.06, 0.25, 0.70) |
| Sequence evolution model | GTR+$\Gamma$ with parameters from Dirichlet (Appendix B) | |
| Estimated gene trees | FastTree ($GTR + \Gamma$) | |

*D1.* This dataset includes 330 replicates and for each replicate, the number of genes were uniformly sampled between 50 and 1000. The number of species were also uniformly sampled between 20 and 200. The birth rate parameter in simulating the species tree is randomly sampled from a log uniform distribution in [10^−7^, 10^−6^], and the death rate is also sampled from a log uniform distribution, bounded from below by 10^−7^ and bounded by the birth rate parameter from above. The population size is sampled from a uniform distribution in [10^5^, 10^6^]. Since sampling multiple individuals is often done for shallow species trees, we sampled a maximum species tree height for each replicate from a log normal distribution with an expected value of 0.5M generations; the number of generations ranges between 0.19M and 1M in 90% of replicates (Fig S1). Moreover, internal branches are extremely short and 90% of them span between 4170 and 382*K* generations.
*D2.* This dataset has three model conditions, each with a fixed maximum number of generations: 0.5M, 1M, or 2M. We simulate 50 replicates of each model condition, all with 200 species and 1000 genes with species birth rate set to 10^−6^ and death rate set to zero (so, a birth-only model). The population size is fixed to 200,000. Each replicate of this dataset also has a single-individual outgroup. To test the effects of sampling, we also create two new versions of D2 where only one or two individuals per species are randomly sub-sampled.

The quartet scores of true species trees given true gene trees, used to measure gene tree discordance due to ILS, indicate that the D1 dataset has extremely high levels of ILS (Fig. 2a). The mean quartet score is 0.496 with standard deviation of 0.127. Note that quartet scores close to 1/3 correspond to gene trees that are random with respect to the species tree. On the D2 dataset, quartet scores range from low (0.50 on average) to relatively high (0.78 on average), depending on the tree height (Fig. 2a).

**Figure 2:**
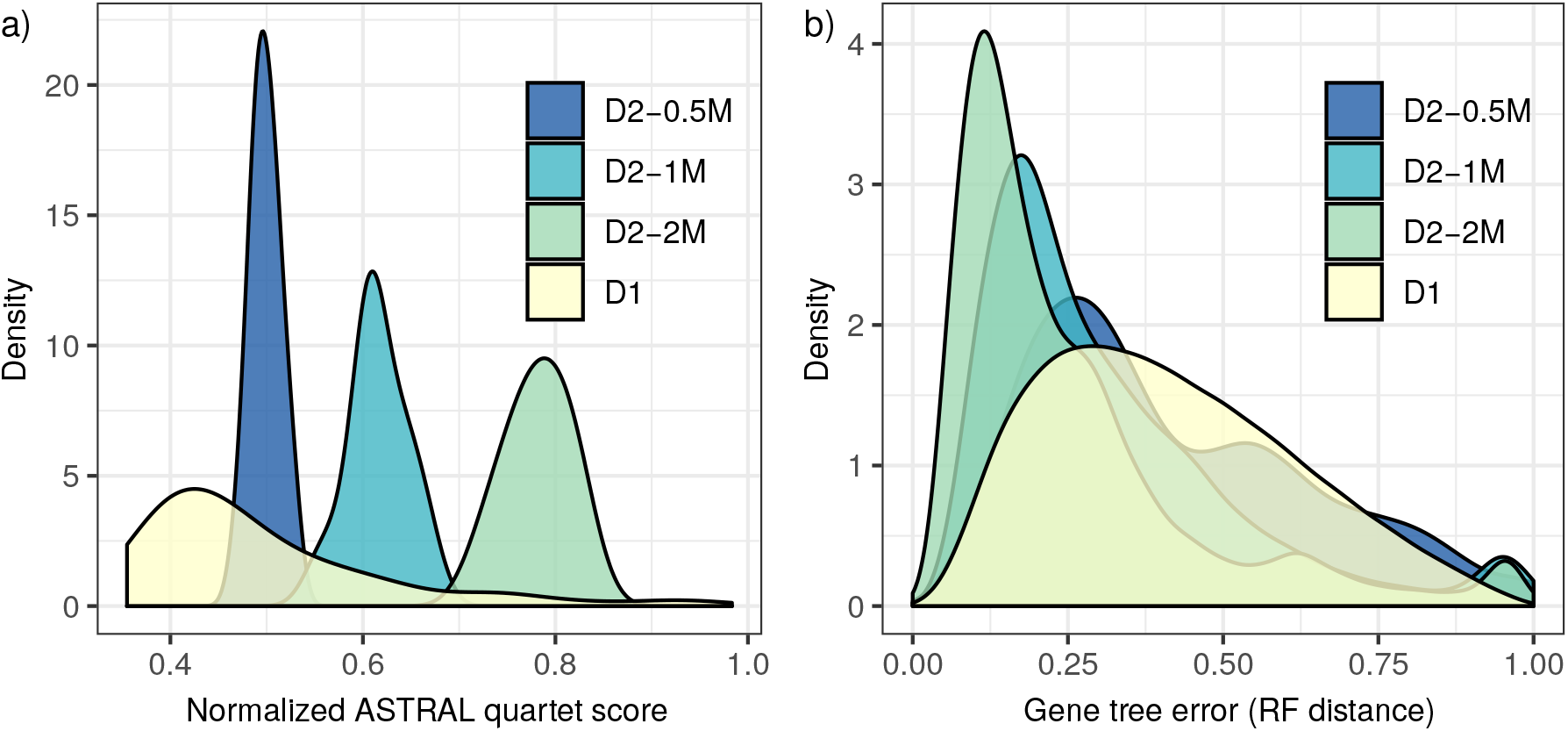
Level of ILS and gene tree error in simulated data. (a) The distributions of the quartet score of the true species tree versus the true gene trees shown as kernel density plots. Colors distinguish datasets D1 (326 replicates) and D2 (50 replicates per model condition defined by tree height set to 0.5M, 1M, or 2M generations). (b) Density plot of normalized Robinson Foulds (RF) distances between estimated and true gene trees, which quantifies gene tree error.

After simulating true gene trees, we use Indelible [47] to evolve nucleotide sequences down the gene trees using the GTR+Γ model of sequence evolution with randomly sampled sequence lengths and mutation parameters. For each gene, we sample the sequence lengths from a log normal distribution with parameters specific to each replicate; the parameters are also drawn randomly from a distribution (described in Appendix B). The empirical average sequence length is 1096 for D1 and 721 for D2. The median gene sequence length is between 406bp and 1803bp in 90% of the 330 replicates of D1 and is between 322bp and 1317bp in 90% of the 150 replicates of D2. The *GTR* + Γ parameters were drawn from Dirichlet distributions used in the ASTRAL-II paper (parameters are estimated from a collection of biological datasets [42]).

Given gene alignments, we then estimate gene trees using FastTree [48] with the GTR+Γ model. The average gene tree estimation error, as measured by the RF distance between true and estimated gene trees, is extremely varied (Fig. 2b). The gene tree error on the D1 dataset is on average 0.4 with standard deviation 0.13, and in 90% of replicates, it ranges between 0.19 and 0.68. On the D2 dataset, the mean gene tree error was 0.42, 0.31, and 0.25 on average for 0.5M, 1M, and 2M model conditions. As expected, shallower trees not only have higher ILS, but also have increased gene tree error (Fig. 2). Both true (simulated) and the inferred gene trees are used as input to summary methods.

### 3.2. Compared Methods

We test several versions of ASTRAL and NJst. When we run ASTRAL-III (version 5.5.4 and version 5.5.9 for branch length and localPP) in the multi-individual mode (i.e., with a mapping file) we refer to the method as ASTRAL-multi. ASTRAL can handle polytomies and previous results indicate that removing very low support branches helps accuracy [44]. Thus, we also test a version of ASTRAL-multi where all the gene trees with branches with support below 5% are contracted and refer to this version as ASTRAL-multi-5%. For D1, we measure support using SH-like supports reported by FastTree. For D2, we compute gene tree bootstrap support with 100 bootstrap replicates. The default ASTRAL-III with each individual treated as a separate species (i.e., with no mapping file) is also tested (ASTRAL-ind). This allows us to investigate whether prespecifying species boundaries helps improving the accuracy of ASTRAL. We gave all ASTRAL runs a maximum of 48 hours of running time. In four out of 330 replicates of D1, either ASTRAL-ind or ASTRAL-multi failed to finish in the allotted time; we exclude these replicates.

NJst (i.e.;, USTAR [41]) is the main existing method capable of handling multiple individuals and *unrooted* gene trees. NJst estimates a distance matrix and uses neighbor joining to construct the species tree. It defines the distance between two species as the average gene tree internode distance or the average number of nodes between the gene copies sampled from the two species across all gene trees. The NJst method is also statistically consistent under the coalescent model after initial errors were fixed [41]. NJst cannot handle polytomies. To address this, we arbitrarily resolve polytomies in gene trees. Note that ML gene trees inferred by FastTree can include polytomies, for example, when multiple sequences are identical in a gene. We also tested removing genes with many polytomies, but this filtering did not help NJst (Fig S2).

### 3.3. Experiments

We study three research questions.

**RQ1:** Does the accuracy of species tree reconstruction improve when multiple individuals are included in a dataset? Does it still improve if the total sequencing effort is kept constant? Using the three models of the D2 dataset with one, two and five individuals, we study effects of the number of individuals per species, either with variable total sequencing effort (i.e., keeping the number of genes fixed) or with fixed sequencing effort. To fix the sequencing effort, we reduce the number of genes such that the product of the number of genes and the total number of individuals is 1000. On the D1 dataset, some replicates have as few as 50 genes; thus, we could not perform these analyses as reducing the number of genes by a factor of 5 would leave us with only 10 genes.
**RQ2:** Is species tree accuracy improved by predefining species boundaries? On the D1 dataset, We compare ASTRAL-ind and ASTRAL-multi. We compute the portion of species that are not monophyletic in ASTRAL-ind and also study the impact of predefined species on the rest of the tree.
**RQ3:** How does the accuracy of ASTRAL compare to alternative methods? We compare the accuracy and running time of ASTRAL-multi and ASTRAL-multi-5% against NJst on both D1 and D2 datasets.

### 3.4. Evaluation Metric

#### Accuracy

To calculate the accuracy of a species tree constructed from input gene trees, we measure the False Negative (FN) rate, defined as the proportion of bipartitions in the true tree that are missing from the estimated tree. Note that since our trees are fully resolved, the FN rate is equal to the Normalized Robinson-Foulds (NRF) distance.

To compare ASTRAL-multi and ASTRAL-ind, we use the extended species tree. Recall that in an extended species tree, each species is replaced by a polytomy containing all individuals of the species. We use the extended tree as the reference tree, compute the FN rate of ASTRAL-ind, and break this FN rate into two components: the FN rate for branches that are terminal in the species tree but are internal in the extended species tree (we call these semi-terminal branches; see Fig 1) and the FN rate for the branches that are internal in both the species tree and the extended species tree (we call these internal branches). The FN rate for the semi-terminal branches gives the percentage of species that have not been recovered as monophyletic in ASTRAL-ind; the FN for semi-terminal branches is zero by construction for ASTRAL-multi.

#### Running time

Running time is measured on the Comet supercomuting cluster with 1 core out of 24 on Intel Xeon E5 CPUs and 5 GB of memory per job.

## 4. Results

### 4.1. RQ1: benefits of sampling multiple individuals

As expected, increasing the number of individuals from one to five gradually reduces the error (Fig. 3a). However, the improvements in accuracy tend to be small and are not statistically significant (*p* = 0.24 according to an ANOVA test with the number of individuals and the number of generations as independent variables); over all 150 replicates of all three conditions of D2, the error is reduced on average from 5.7% to 5.2% when going from a single individual to five. Contrary to our expectations, the impact of the number of generations (tree depth) on the effectiveness of increasing the number of individuals was also not significant (*p* = 0.85).

**Figure 3:**
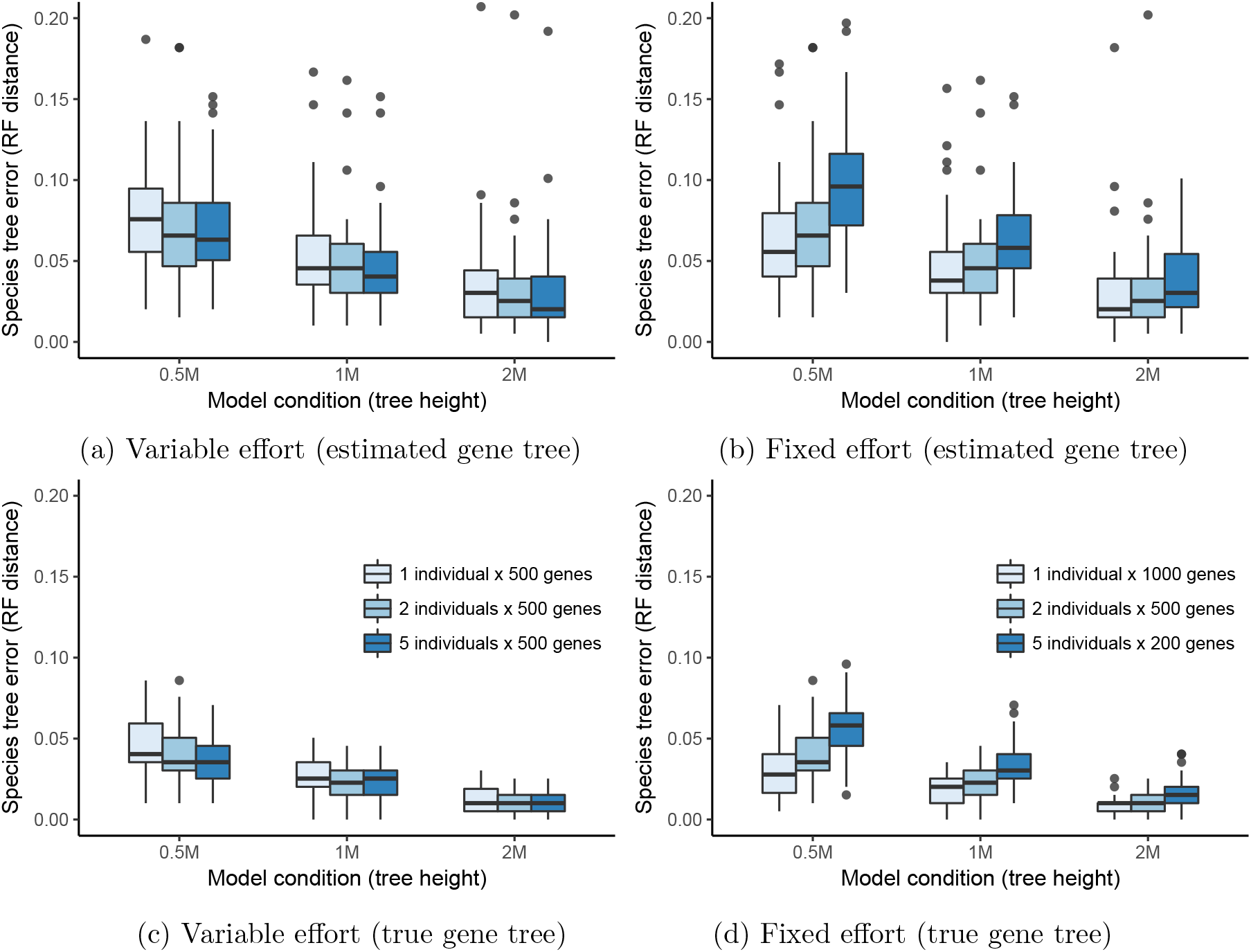
Impacts of increased individual sampling. The Robinson Foulds (RF) distance of the ASTRAL-multi species tree to the true tree is shown for three model conditions (x-axis) of the D2 dataset with (a,b) estimated gene trees and (c,d) true gene trees as input and with (a,c) variable sequencing effort or (b,d) fixed effort. Boxplots show the median (bars), the interquartile range (boxes), and outliers (points) defined as points above/below whiskers, which extend up to 1.5 times the height of the interquartile range on each side.

When we fix the total sequencing effort by reducing the number of genes as we increase the number of species, it becomes clear that sequencing more individuals is not nearly as effective as sequencing more genes (Fig. 3b). Thus, the error with 1000 genes and a single individual per species (4.7% on average overall) is less than the error with 200 genes and five individuals (7.0% on average overall). The same pattern is observed for all three model conditions.

Unlike estimated gene trees, when true gene trees are used, improvements in the accuracy are substantial (Fig. 3c), especially for the shallow model condition (0.5M generations). With true gene trees, improvements in accuracy with variable effort are indeed statistically significant (*p* = 0.017). Nevertheless, even with true gene trees, fixing the effort shows that having more genes is more effective than increasing the number of individuals (Fig. 3d).

### 4.2. RQ2: benefits of defining species boundaries

When we run ASTRAL-ind (i.e, without defining species) on the D1 dataset, we observe that the FN error of the resulting species tree is somewhat higher than when the mapping is predefined (Table 2). Focusing on the FN error rate of semi-terminal branches shows that close to 9% of the species are not recovered as monophyletic when the mapping is not known in advance. Moreover, even focusing on internal branches, ASTRAL-ind has slightly higher error (8.3%) than ASTRAL-multi (7.8%), which indicates that providing species boundaries can also improve the accuracy of detecting the relationships among species. Nevertheless, we caution that the improvements, while statistically significant (*p* < 10^−5^ according to a paired t-test) are not large.

**Table 2:**
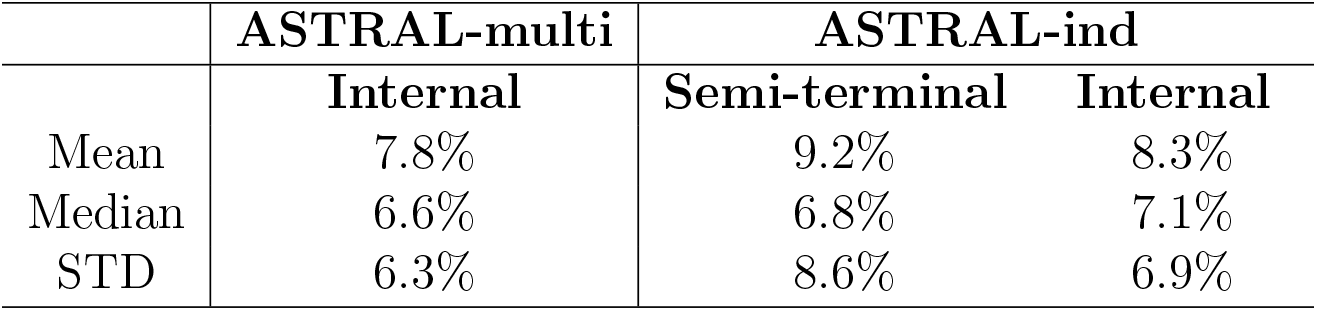
The species tree error of ASTRAL-multi and ASTRAL-ind on the D1 dataset. For ASTRAL-multi, we show the average (326 replicates) Robinson Foulds (RF) distance to the true species tree, which is identical to False Negative (FN) rate. For ASTRAL-ind, we compare the inferred tree to the extended species tree, and show the FN rate, divided into two categories: semi-terminal branches (i.e., those corresponding to species) and internal branches (i.e., all other branches).

|  | ASTRAL-multi | ASTRAL-ind |  |
| --- | --- | --- | --- |
|  | Internal | Semi-terminal | Internal |
| Mean | 7.8% | 9.2% | 8.3% |
| Median | 6.6% | 6.8% | 7.1% |
| STD | 6.3% | 8.6% | 6.9% |

The error of ASTRAL-ind in recovering species as monophyletic is greatly impacted by the amount of ILS (Fig. 4a). For the highest levels of ILS, the error for semi-terminal branches can be as high as 20%, but this gradually reduces to 2% as the quartet score increases and the amount of ILS decreases. Similarly, the error in the internal branches of ASTRAL-multi is lower than ASTRAL-ind mostly for high levels of ILS and less so as the amount of ILS decreases (Fig. 4a). Interestingly, the wrong semi-terminal branches have low localPP; removing branches with 0.95 localPP or lower make ASTRAL-ind trees compatible with the monophyly of almost all the species (Fig. 4b). Another benefit of prespecifying species boundaries is that ASTRAL-multi runs around 23% faster that ASTRAL-ind on the D1 dataset.

**Figure 4:**
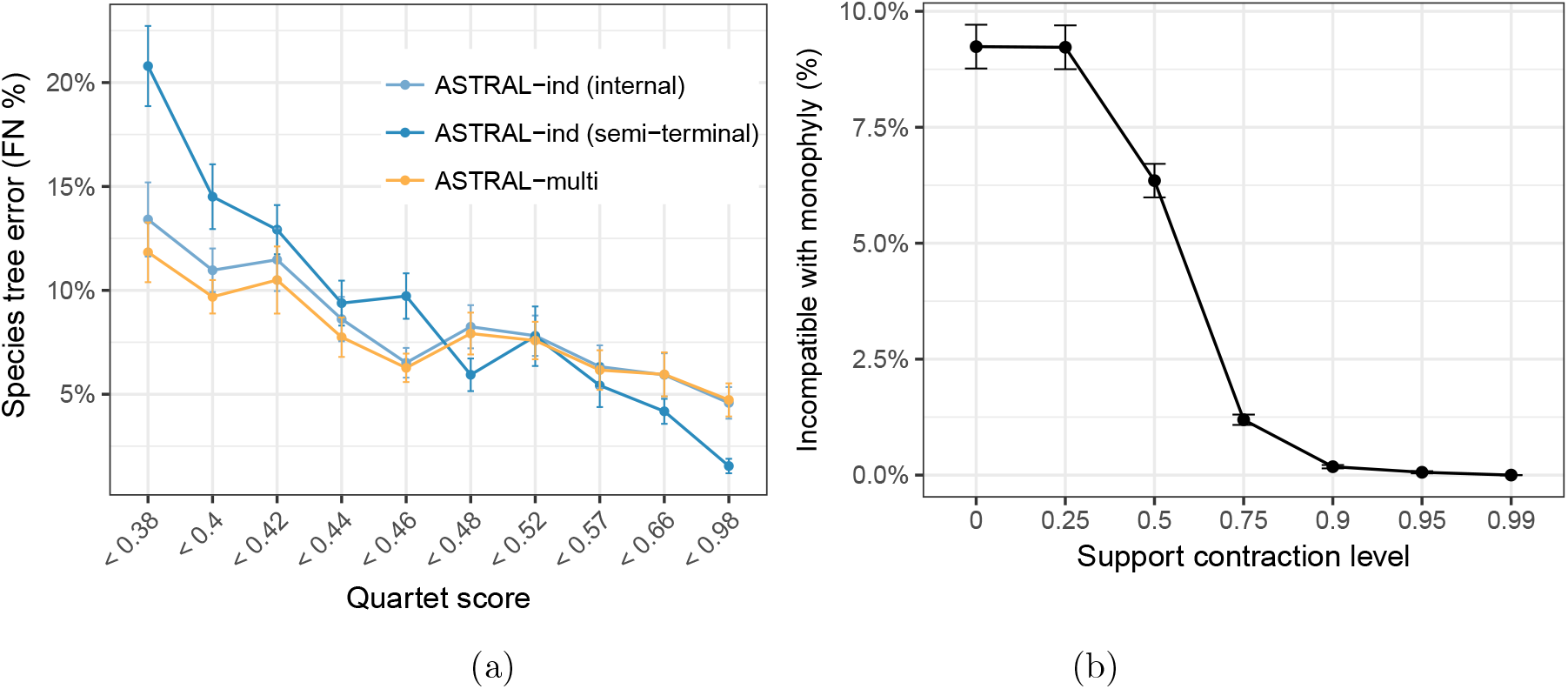
Impact of predefining species. (a) We show the proportion of branches of the true species tree missing in the trees inferred by ASTRAL-multi and ASTRAL-ind (without known species boundaries) on the D1 dataset (FN rate). For ASTRAL-ind, the reference tree is the extended species tree and the error is calculated separately for semi-terminal and internal branches. The 326 replicates are divided into deciles based on their level of ILS, measured by the quartet score of the true species tree (x-axis). The mean and standard error of FN are shown. (b) The percentage of species that are not compatible with monophyly in species trees constructed by ASTRAL-ind on D1 after branches below a level of localPP support (x-axis) are contracted.

### 4.3. RQ3: method comparison

We now compare the error rate of ASTRAL-multi and ASTRAL-multi-5% to NJst. On the D1 dataset, methods performed similarly in terms of accuracy (Fig. S3). In terms of running time, ASTRAL-multi is about three times faster than NJst (Table S1). In the rest of this section, we will focus on on the D2 dataset.

#### Accuracy

On the D2 dataset with five individuals per species, ASTRAL-multi-5% is in most cases, but not always, better than ASTRAL-multi (Fig. 5). The comparison between ASTRAL-multi-5% and NJst depends on the dataset. With very shallow trees (0.5M condition), ASTRAL-multi-5% and NJst are essentially tied, with a small advantage to NJst but the differences are not statistically significant (*p* = 0.30 according to a two-way ANOVA test with the choice of the method and the number of genes as independent variables). However, with the longer trees (1M and 2M conditions), ASTRAL-multi-5% is significantly better than NJst (*p* = 0.012), and the extent of the improvement is not significantly impacted by the number of genes (*p* = 0.81).

**Figure 5:**
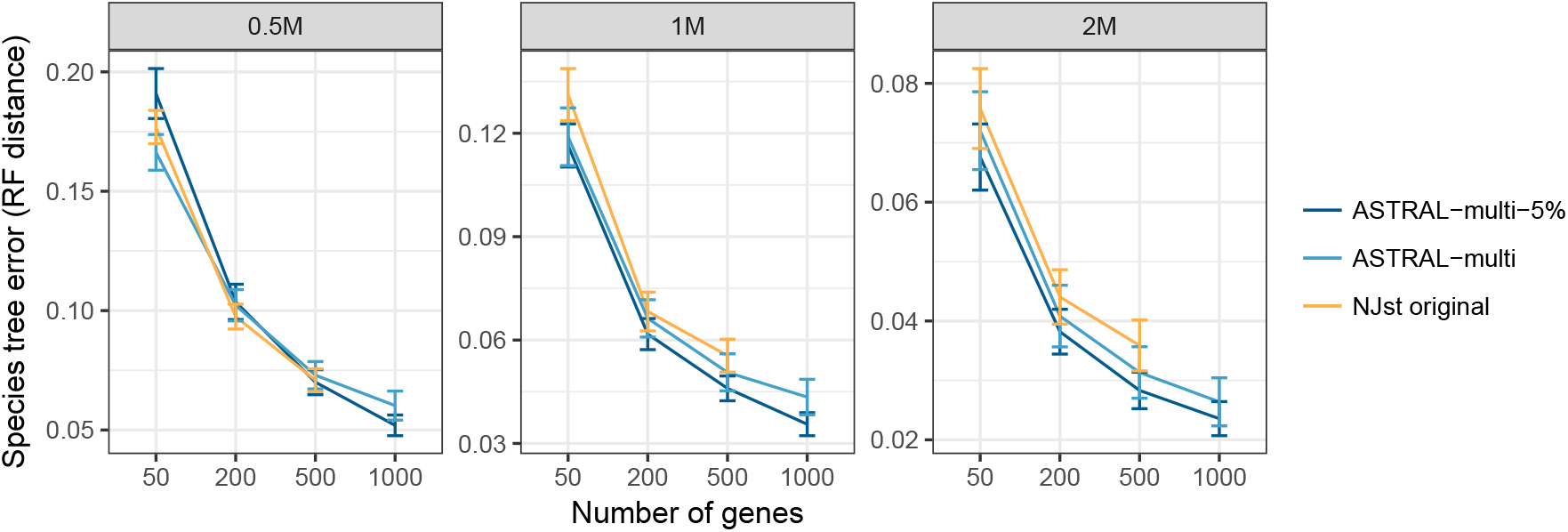
The error rate of ASTRAL-multi, ASTRAL-multi-5% and NJst on three models of dataset D2 with four levels of number of genes which are 50, 200, 500 and 1000. Mean and standard error of species tree error is shown over 50 replicates per condition.

Overall patterns are as expected. Increasing the number of genes reduces the error while reducing the number of generations increases error. Not only shallow trees are harder to resolve, but also, smaller differences are observed between alternative methods for shallow trees. This could perhaps indicate that high gene tree error (Fig S1b) in shallow trees reduces the input quality and erases any potential differences between various summary methods.

#### Running Times

We compare the running time of ASTRAL-multi to NJst on D1 which has replicates with different input size (number of genes multiplied by number of leaves) from about 2000 to 16000. ASTRAL is substantially faster than NJst even running with contracted data (Fig S4). Given two days of running time, ASTRAL-multi and ASTRAL-multi-5% were able to finish on 327 and 324 replicates, respectively whereas NJst was able to finish on 313 replicates. A faster implementation of NJst called ASTRID [49] cannot currently handle multi-individual inputs.

## 5. Discussion

We introduced a multi-allele version of ASTRAL and on two large-scale simulated datasets we demonstrated its accuracy and running time efficiency. We saw that predefining the species boundaries can improve the accuracy. Sampling multiple individuals did not seem to help the accuracy.

### Effectiveness of individual sampling

Unsurprisingly, increasing the number of individuals somewhat helped accuracy, but in the presence of gene tree estimation error, the improvements were marginal. Moreover, compared to increasing the number of genes, increasing the number of individuals did not seem an efficient use of resources. It can be argued that our way of fixing “effort” by controlling the product of the number of genes and the number of individuals is naive, as the cost of increasing loci versus individuals may differ. The exact relative costs will depend on the sequencing technology, sample collection, and many other factors beyond the scope of our work. Nevertheless, our results point to limited effectiveness of sampling more individuals.

It is perhaps surprising that even with variable sequencing effort, the reduction in the error is small as we increase the numbers of individuals. Our results are somewhat contrary to some previous simulation studies [12, 33, 32] that indicate that sampling more individuals is beneficial for shallow trees but agrees with others [40]. We stress that many of our simulations, especially D1, were on extremely shallow trees. For example, on the D1 dataset, our trees on average included 110 species generated in about half a million generations. Thus, the species tree branch lengths were extremely short; 70% of branches were a hundred thousand generations or shorter and 47% had a length ≤ 0.1 in coalescent units. Thus, our simulations were specifically designed to test conditions where multiple individuals may help according to previous reports, making it even more interesting that no strong pattern of improvement was observed. Our simulations differed from previous works in the level of gene tree error and also in the number of genes. Our gene trees have high levels of error (Fig. S1). It may be that gene tree estimation error for shallow trees reduces the value of having multiple individuals because the extra noise introduced by tree estimation weakens the signal. Consistent with this explanation, with true gene trees, using more individuals did result in improvements. Another difference between our study and previous studies is that we use hundreds of gene trees whereas previous studies include at most 50 gene trees [12, 33, 32, 40]. Another explanation is that with enough loci, the impact of having more individuals diminishes. We believe that with the current sequencing technology, testing methods in the presence of hundreds of loci is more relevant than tens of loci.

### Errors in species delimitation

In all our analyses, the predefined species boundaries were perfectly correct. In practice, the boundaries defined *a priori* may or may not be correct. Introducing error in species identification may erase some of the benefits of using the multi-allele version of ASTRAL. Two solutions can be employed in practice. Helpfully, our simulations showed that contracting low support branches made all species compatible with monophyly (Fig. 4b). Thus, to find mistakes in the species mapping, the dataset can be analyzed in both modes: with and without species boundaries. When strong localPP is found for the lack of monophyly of some species, the analyst can reconsider the delimitation. Moreover, the multiallele version of ASTRAL can produce branch length and localPP for the terminal branches of the species tree. In our simulations, terminal branches of the species tree generally had high support (Fig. 6c). For example, 75% of them had a localPP of 1.0 and close to 90% had a localPP of 0.75 or higher. When terminal branches have very low support, the species definition should be questioned.

**Figure 6:**
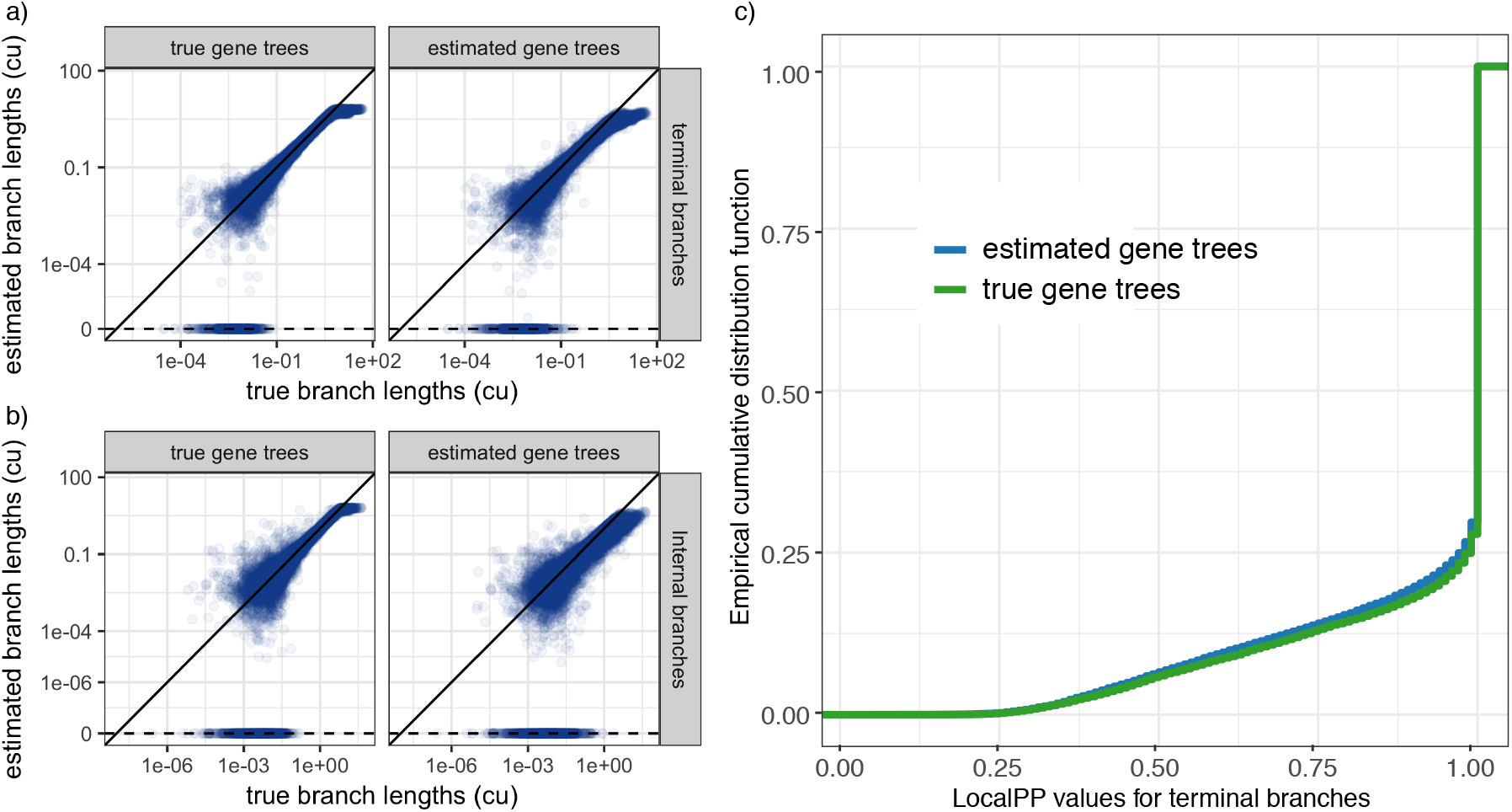
The accuracy of branch length and localPP support. (a,b) On the D1 dataset, we estimate the length of (a) terminal and (b) internal branches of the true species trees using both true and estimated gene trees in coalescent unit (cu). We show the true branch length (x-axis) versus the estimated branch length (y-axis), both in the log scale. (c) The empirical cumulative distribution function of the localPP support for terminal branches of the true species trees for D1 using estimated and true gene trees. Close to 75% of the terminal branches have the maximum localPP support.

### Branch length accuracy

ASTRAL-multi can compute both terminal and internal branches. In our simulations, when true gene trees were used, terminal and internal branches lengths were both relatively accurate (Figs. 6ab and S5). However, the accuracy reduced substantially with estimated gene trees. Errors of close to even an order of magnitude were observed in estimated branches. This pattern, which is consistent with older results [45], indicates that in the presence of gene tree error, branch lengths should be considered with caution. Interesting, we note that terminal branches seemed to have less error than internal branches (Fig. 6ab). Finally, using multiple individuals instead of one individual resulted in a small but consistent improvement in internal branch lengths, but once again, increasing the number of genes was more effective (Fig. S5).

### Predictors of accuracy

The heterogeneity of the D1 dataset enables us to look for parameters that impact the accuracy of the ASTRAL species tree (Fig 7). We observed a strong dependence between the species tree error and the number of genes, the tree depth, the population size, the amount of true discordance measured by the quartet score of the true species tree versus true gene trees, and the average gene tree error. The only factor we studied that did not have a clear impact on the accuracy was the number of species. All factors other than gene tree error had a similar impact on the species tee accuracy if the true gene trees were used (Fig. S6). As expected, main predictors of the accuracy seemed to be the number of genes, the amount of true discordance, and the gene tree estimation error (Fig 7). These three factors combined in a simple linear model could explain 40% of the variation in the species tree error. The true discordance measured by the quartet score, when log-transformed, seemed to correlate close to linearly with the species tree error. The other two factors had more intricate patterns of impact.

**Figure 7:**
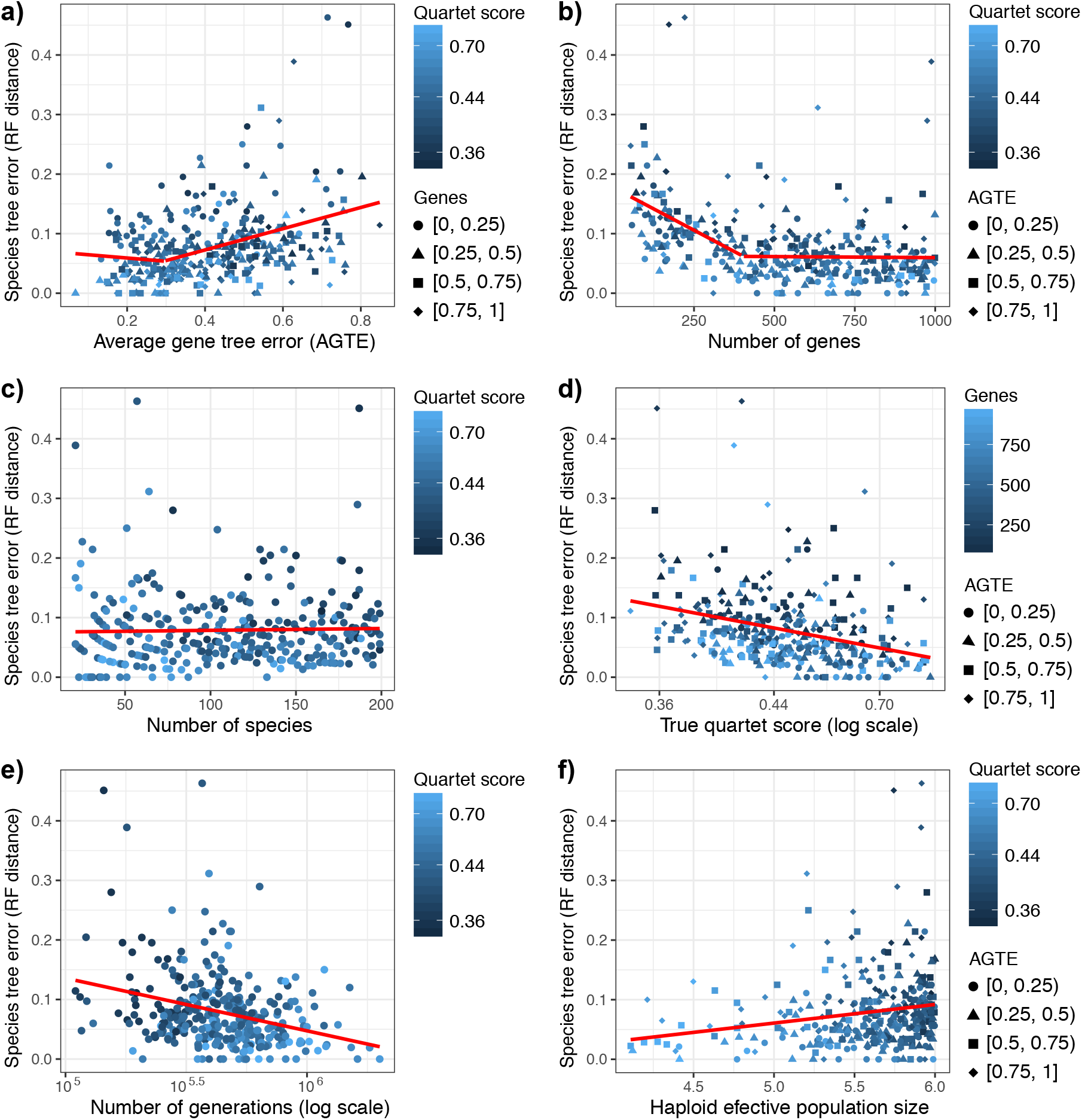
The impact of simulation parameters on the ASTRAL-multi species trees. We show Robinson Foulds (RF) distance between the ASTRAL-multi species tree and the true species tree (y-axis) versus (a) average gene tree error (AGTE), (b) the number of genes, (c) the number of species, (d) the quartet score of the true species tree based on true gene trees (showing the ILS level), (e) the number of generations in log scale, and (f) haploid effective population size. Shades of blue show the quartet score in all panels except (d) where they show the number of genes. Point shapes distinguish quartiles of AGTE (b, d, f) or the number of genes (a). A linear model (red line) is fitted to the data; in (a,b), a two-segment liner model is fitted, with breakpoints determined using the bootstrap fitting [50] algorithm.

Increasing the number of genes to ≈400 had a strong effect on the species tree accuracy (correlation coefficient: −0.44 in a linear fit). Beyond 400 genes, the impacts on accuracy were diminished (slope: ≈ 0; correlation coefficient: −0.06). When the number of genes was 800 or more, it was rare (5 replicates) that the species tree error was above 0.1 NRF, and when it did, the gene tree error was typically high. Gene tree error also had non-linear effects. When the average gene tree estimation error estimated by NRF was below 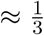, it had only a small impact on the species tree error, but beyond this threshold, it strongly impacted the species tree accuracy. However, it should be noted that even when the gene tree error is 0.6 NRF or higher, there are two replicates that have a large number of genes and a relatively low true discordance, and in both cases, ASTRAL has close to perfect accuracy. Conversely, when gene tree error is as low as 0.34, a replicate that has only 85 genes and very high discordance (quartet score= 0.37) had a relatively high species tree error (0.2 NRF error). Thus, to predict the accuracy, all three factors have to be considered.

### Adequacy of the set X

To test whether the set *X*, the constrained set of bipartitions searched by ASTRAL, is sufficiently large, we ran ASTRAL with bipartitions of the true species tree forced to be in *X*. When this enforcement improves the quartet score or the accuracy, we can infer *X* is not sufficiently large; on the other hand, if making sure *X* includes all bipartitions of the true tree does not improve the score or the accuracy, then, further expanding *X* is unlikely to improve the accuracy. In our simulations, the median species tree error and quartet scores remained constant with the addition of true bipartitions and the average improved by only 0.0004 for accuracy and by 3 × 10^−6^ for the quartet score (Table S2). Thus, we conclude that the search space is sufficiently large. Moreover, as expected, |*X*| is a function of *m* and *k* and just like single individual datasets, we observe that |*X*| grows linearly with *mk* (Fig. S7).

### Older versions of ASTRAL-multi

Even though we are describing the multi version of AS-TRAL for the first time, previous versions of our algorithm have been made available publicly previously. Some published papers have already used those older versions to obtain their species trees [51, 52, 53]. Compared to previous versions (< 5.0.0), the new version is expected to be faster. It is not clear to us whether there are cases where the new version will give better quartet scores. We reanalyzed the 498-locus dataset of 112 species and 163 individuals from the genus Protea [51] using the latest version of ASTRAL-multi and compared the results to version 4.7.9 used by the original paper. The new version produces an identical tree, but the search space *X* has now reduced from 208505 clusters to 64183 cluster, resulting in a 4X reduction of the running time.

## Supplementary Material

### Appendix A. Supplementary figures

**Table S1:** Running times statistics of running ASTRAL and NJst on D1 in hours measured on the same clusters (Comet). 313 out of 330 replicates are reported here since NJst failed to finish on 17 of them in 48 hours. Also ASTRAL-multi could not finish 3 of them and ASTRAL-multi-5% fails on 6 of them.

|  | ASTRAL-multi | ASTRAL-multi-5% | NJst |
| --- | --- | --- | --- |
| Mean | 3.745 | 4.124 | 10.658 |
| Median | 1.528 | 1.571 | 5.737 |
| STD | 5.227 | 5.995 | 11.400 |

**Table S2:** FN and quartet score difference of the species tree constructed from the set of gene trees of D1, with and without adding clusters of true species tree. The values shown are calculated such that positive FN diff means how much adding clusters of true species tree helps and negative quartet score diff shows how much quartet score has increased with adding true species tree bipartitions to search space.

|  | Quartet score diff | FN diff |
| --- | --- | --- |
| Mean | -2.98673e-06 | 0.0004 |
| Median | 0.0 | 0.0 |
| STD | 2.00276e-05 | 0.0066 |

#### Algorithm S1 Computing similarity matrix

*getSimilarity* is defined in [42] and gives the similarity matrix of the leaves of its input gene trees, labeled with individuals names.

~~~
**function** GETSPECIESSIMILARITY(*𝒢*)
   *GS* ← *getSimilarity*(*𝒢*)
   *S* ← *Zeros*(*n* × *n*)
   **for** *i* ∈ *Rows*(*GS*) **do
      for** *j* ∈ *ols*(*GS*) **do**
              *S*[*i, j*]+ = *GS*[*s*(*i*)*, s*(*j*)]
   *D*[*i*] = |{*j*|*s*(*j*) ∈ *i*}|
   *S*[*i, j*] = *S*[*i, j*]/*D*[*i*] × *D*[*j*]
~~~

**Figure S1:**
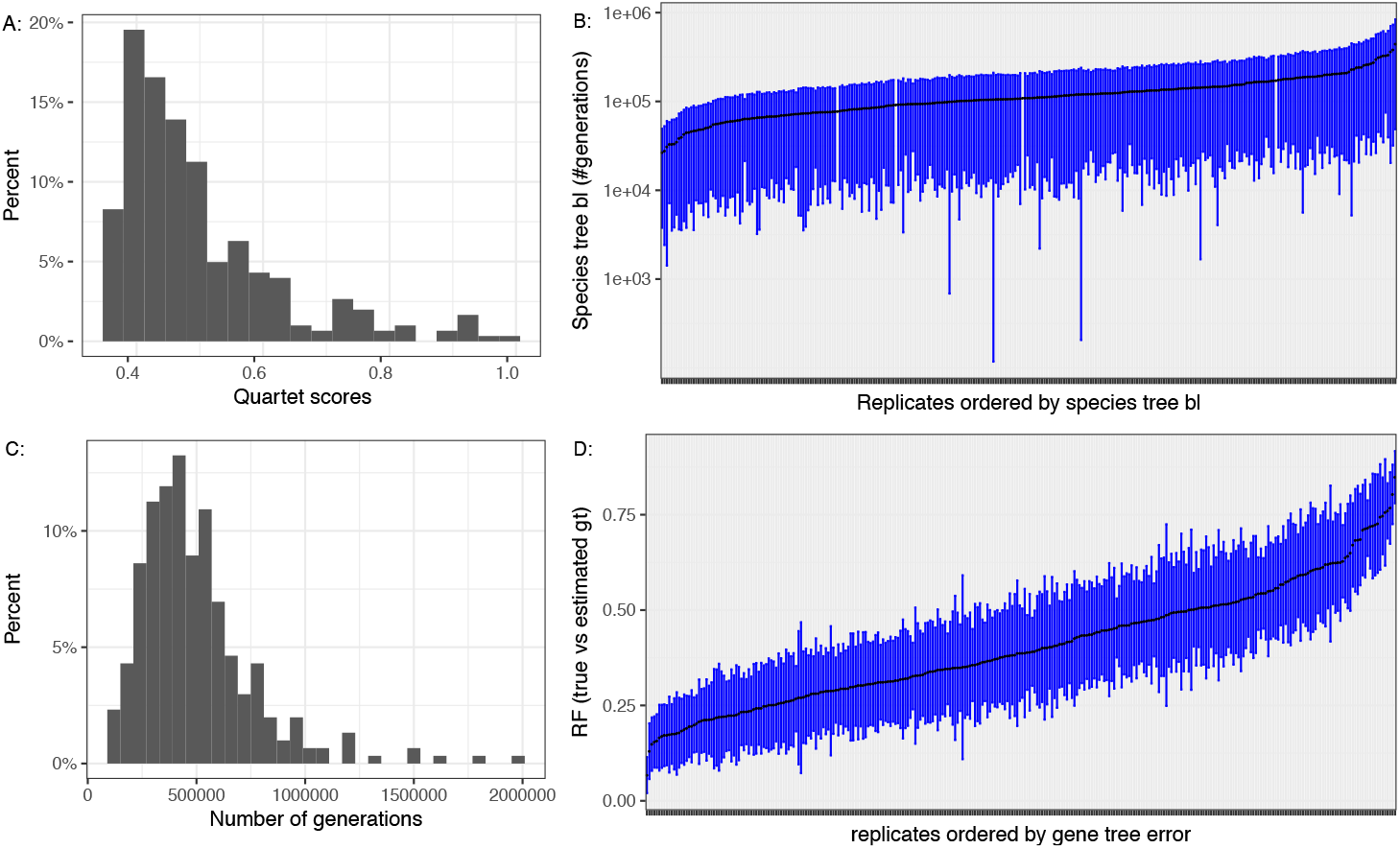
Simulation properties for D1. A) The empirical histogram of the quartet scores of ture species tree using true gene trees on D1. There are a few replicates with high ASTRAL quartet score. In 90% of replicates the quartet score is between 0.37 and 0.76 and the average ASTRAL quartet score is 0.5. B) The average (points) and standard deviation (bars) of species tree branch lengths measured in the number generations (log scale). The number of generations between speciation events ranges between 10,000 to 1000,000 in most cases. C) The empirical histogram of the tree height measured in the number of generations. D) The average (points) and standard deviation (bars) of the RF distance between true and estimated gene trees.

**Figure S2:**
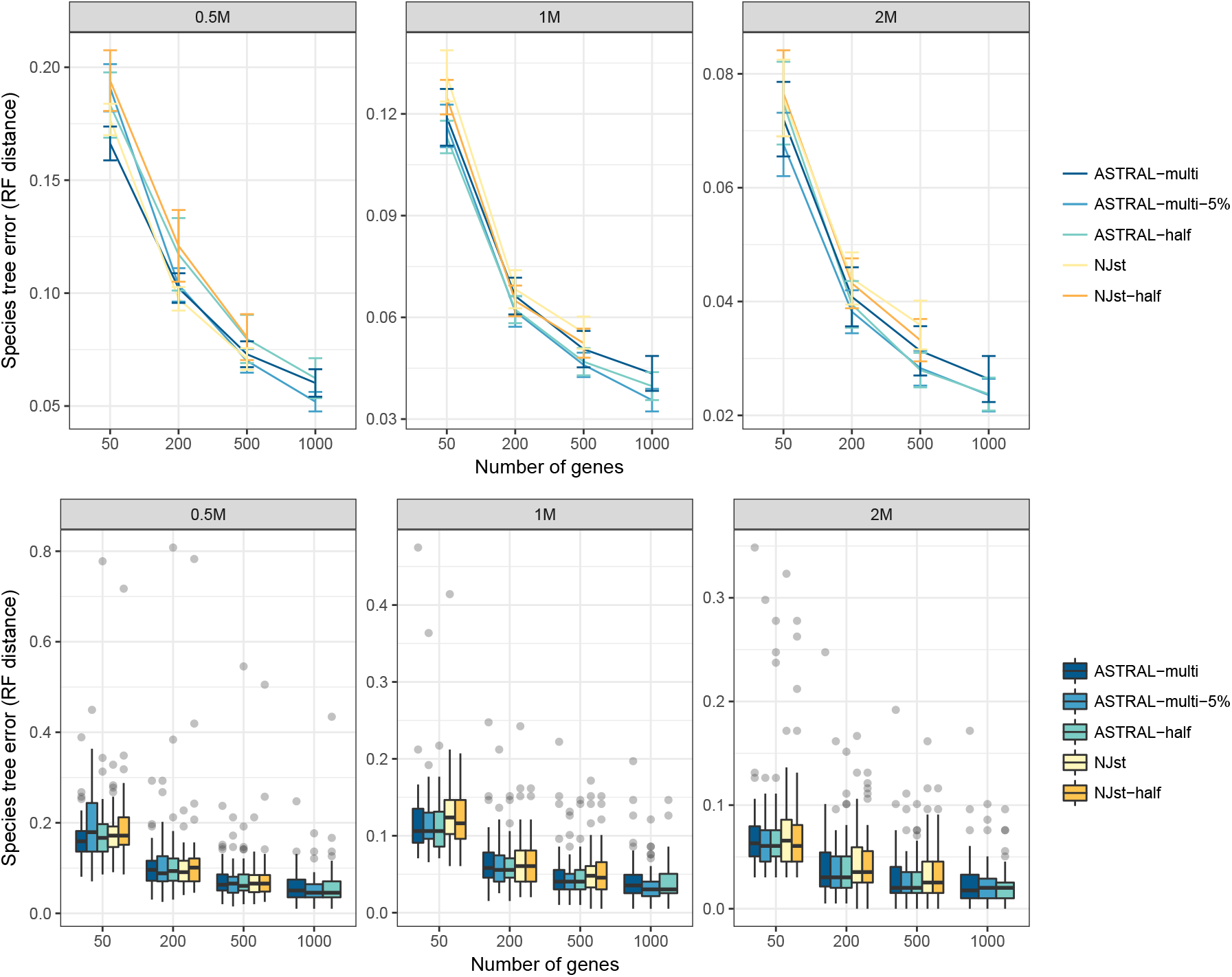
Species tree error (RF distance) of species tree constructed from estimated gene trees with different versions for ASTRAL and NJst on the D2 dataset. Top: mean and standard error over all 50 replicates for each model condition; Bottom: The bolxplots over the 50 replicates. ASTRAL-half is like ASTRAL-multi but genes with many polytomies are removed from the input set. Genes with many polytomies are defined to be those where the number of internal branches is at most half of the maximum possible number of internal branches. Similarly, for NJst-half, these genes are removed. This filtering does not produce a clear improvement in NJst or ASTRAL.

**Figure S3:**
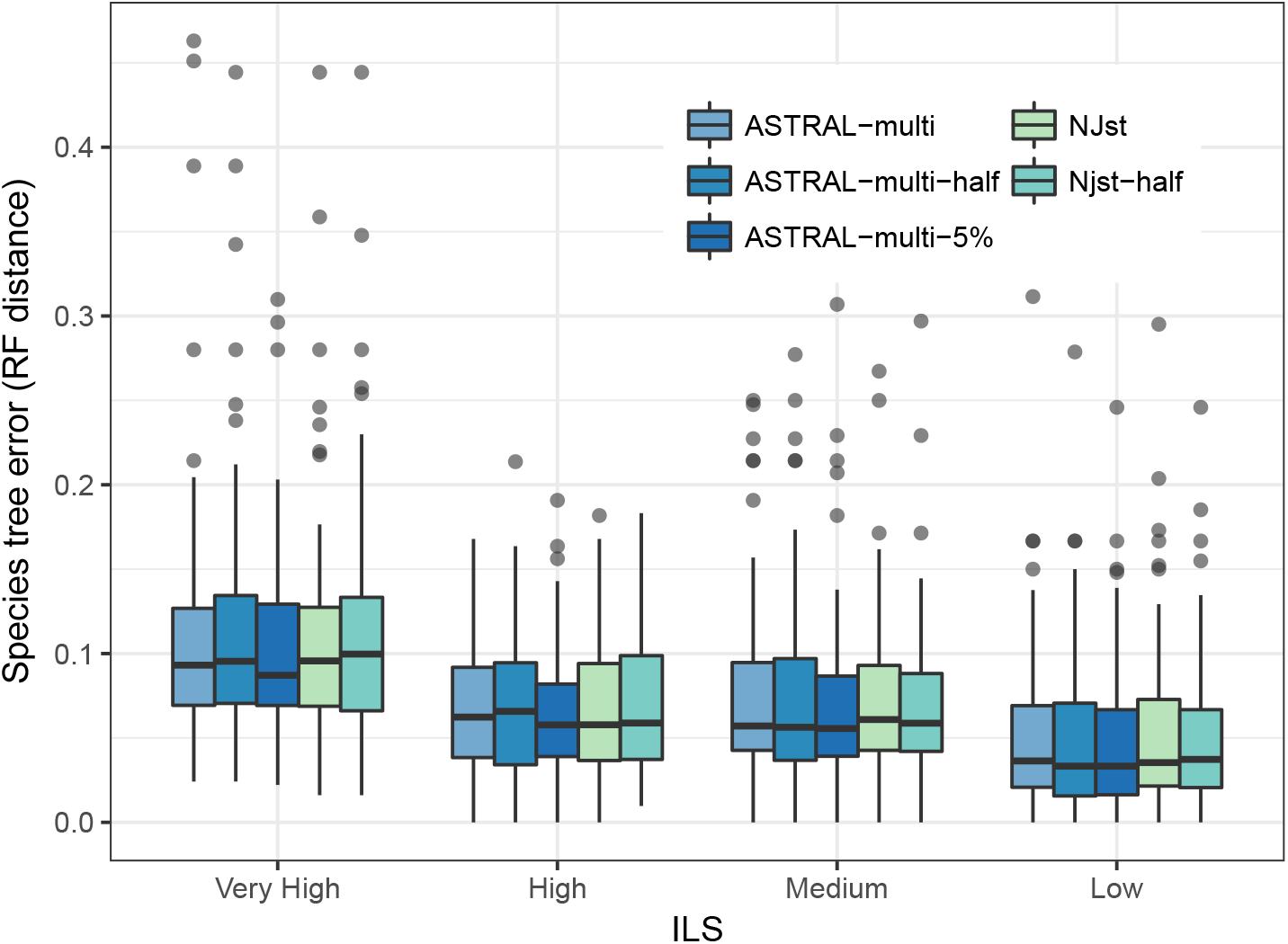
Species tree error of three version of ASTRAL versus two version of NJst run on the D1 dataset. The 326 replicates are divided into quantiles according to their ILS level, as measured by the quartet score (bottom). ASTRAL-multi-half is running ASTRAL-multi but genes with many polytomies are removed from the input set. Genes with many polytomies are defined to be those where the number of internal branches is at most half of the maximum possible number of internal branches. Similarly, for NJst-half, these genes are removed. On this dataset, all methods perform similarly.

**Figure S4:**
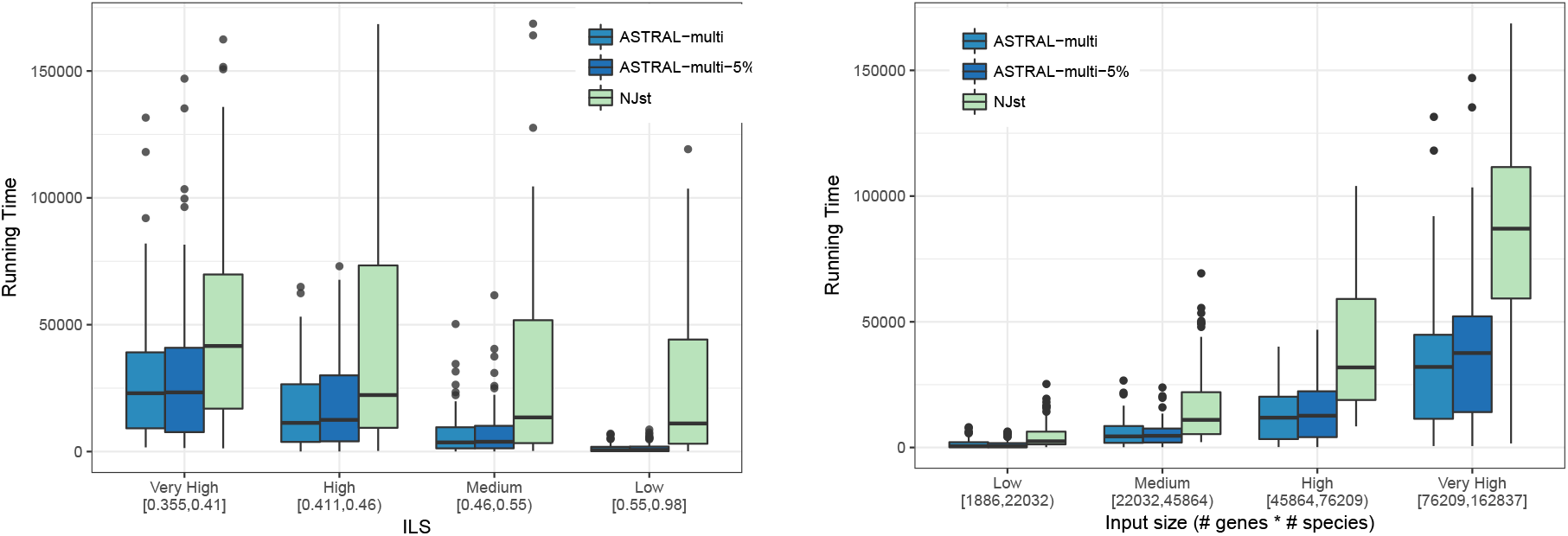
Running time comparison between ASTRAL-multi, ASTRAL-multi-5% and NJst in seconds with respect to ILS level and input size defined as number of genes multiplied by number of species. ASTRAL is faster than NJst in all conditions. The running time of the ASTRAL-Multi increases with increased ILS or with larger inputs.

**Figure S5:**
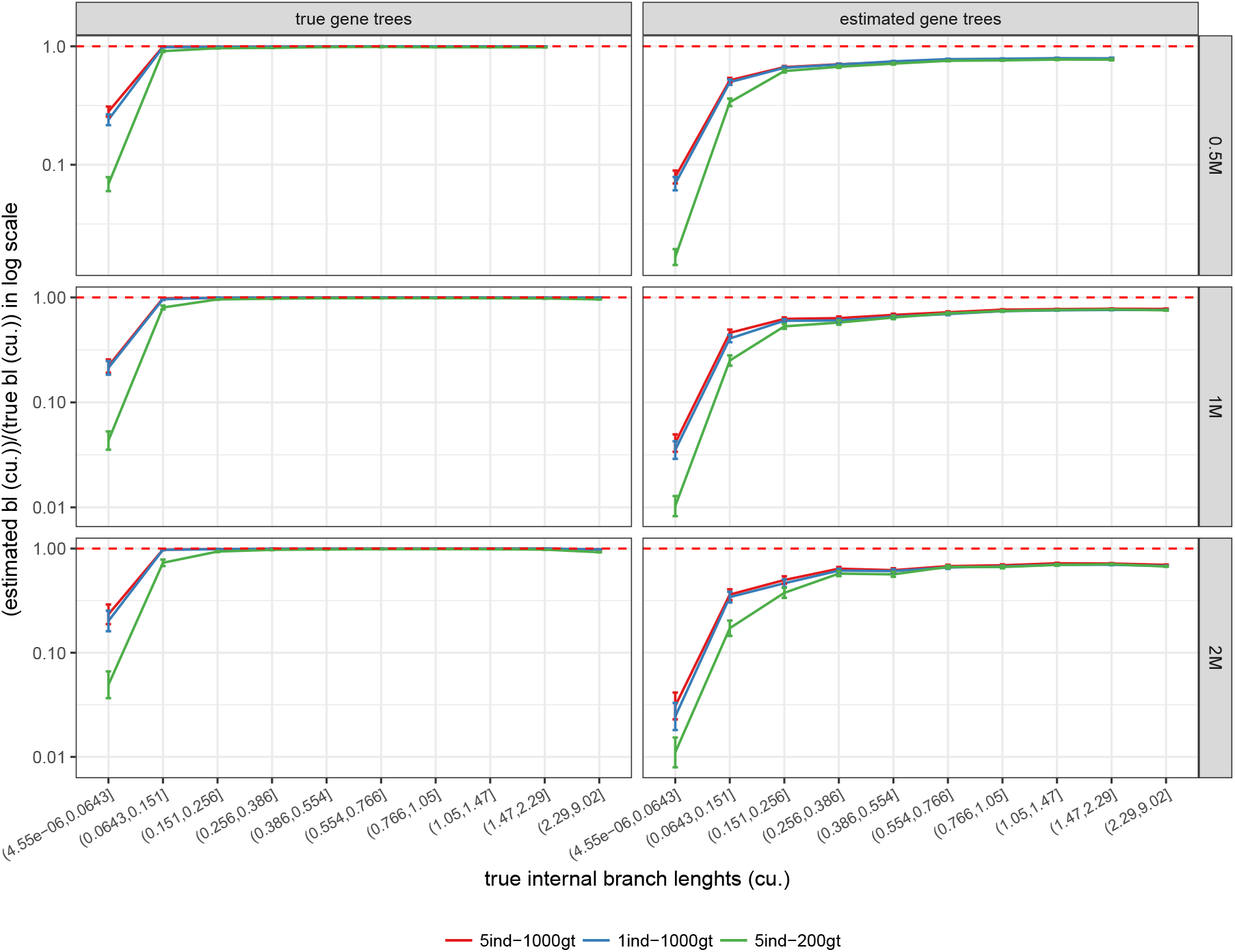
Accuracy of internal branch lengths for D2 using true species trees and estimated and true gene trees. x-axis: deciles of true branch lengths, y-axis: average ratio of estimated internal branch lengths to true internal branch lengths in log scales. Dotted red line indicates ratio of 1, which indicates perfect accuracy. solid red line: 5 individuals and 1000 gene trees, solid blue line: 1 individual and 1000 genes, solid green line: 5 individuals and 200 gene trees. Increasing the number of individuals at the expense of the number of genes reduces the accuracy of branch lengths.

**Figure S6:**
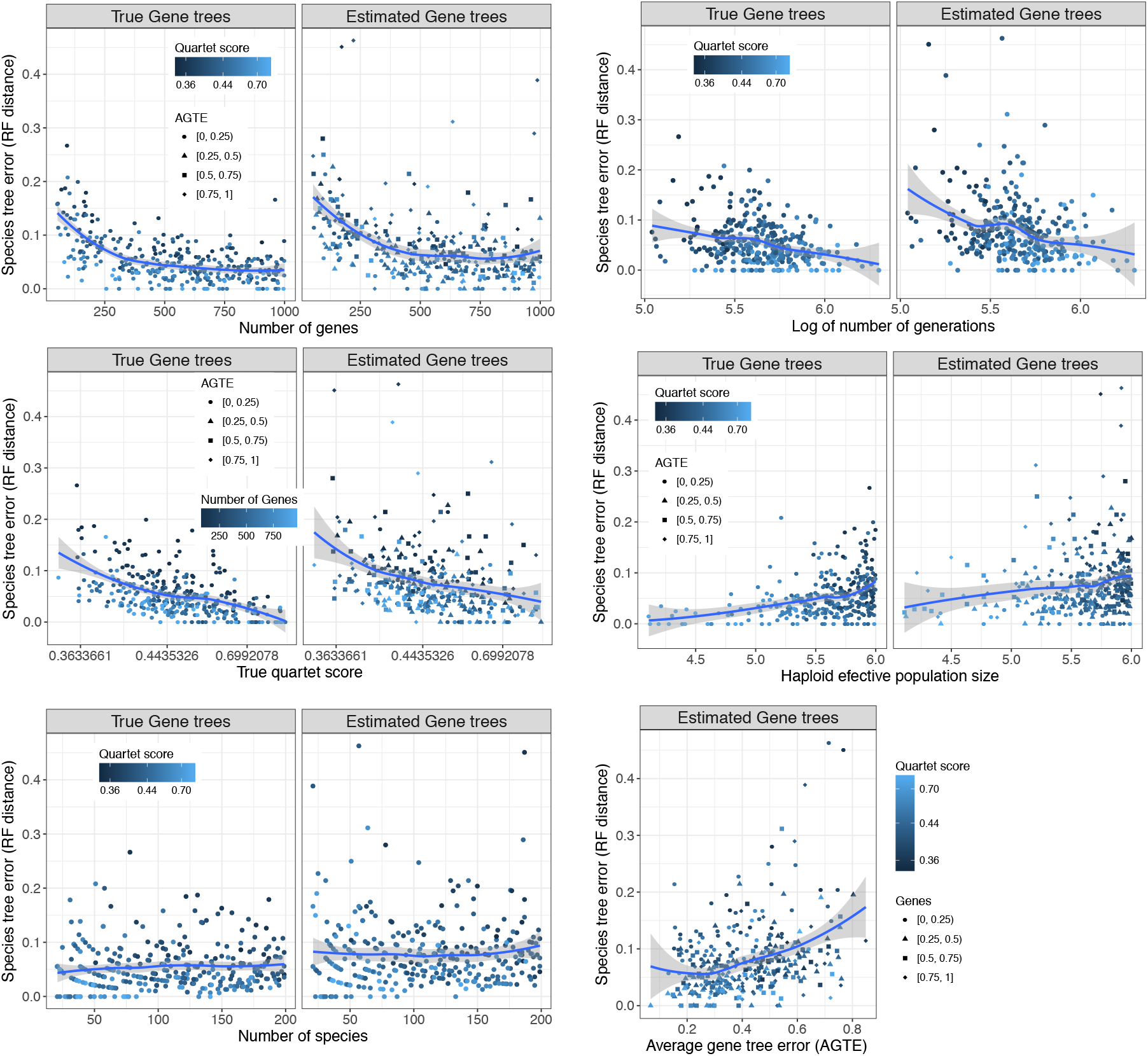
FN error of species trees constructed from estimated gene trees with respect to number of genes, log number of generations, true quartet score, haploid effective population size, number of leaves and average gene tree error (AGTE).

**Figure S7:**
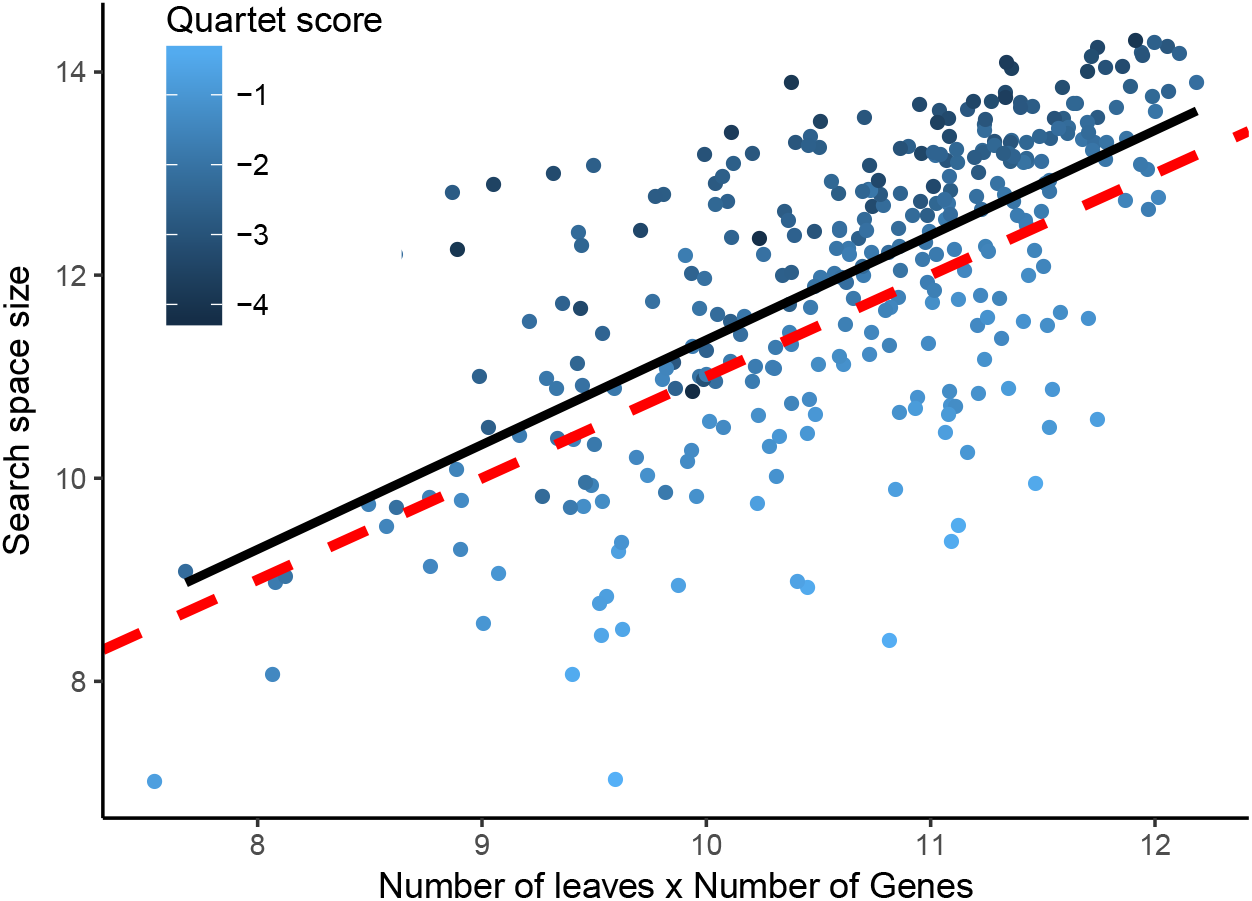
Number of clusters in the search space of ASTRAL-multi versus the size of the dataset on the D1 dataset. The search space size is quantified as ln(|*X*|) and the dataset set size is ln(*mk*) where *m* is the number of leaves and *k* is the number of genes. The red dotted line: ln |*X*| = *α* + ln *mk*. The black line: a line fitted to the data. Colors: (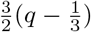) where *q* is the quartet score; lighter values show higher quartet scores and thus, less ILS. Increasing the dataset size linearly increases the search space size (because the black line and the red dotted lines have similar slopes). The quartet scores is a large driver of the variation for each dataset size.

### Appendix B. Simulation procedure

In order to generate D1 we used Simphy [46] with the following exact command

~~~
simphy −rs 330 − rl u : 5 0, 1000 −rg 1 −sb lu : 0 . 000000 1, 0. 000001
−sd lu : 0 . 0000001, sb −s t l n : 13, 0.5 − sl u:20,200 − si f:5
−sp u : 10000, 1000000 −su ln : − 17 . 27461, 0 . 6931472 −hs ln :1.5, 1
−hl ln:1.551533, 0.6931472 −hg ln : 1.4, 1
−cs 9644 –v 3 −o tre −ot 0 −op 1 −od 1
~~~

For D2 with 0.5M generations we used

~~~
simphy −rs 50 − rl U1000, 1000 −rg 1 −st U1000000, 1000000 − si U5, 5
− sl U200, 200 −sb U0 . 000001, 0 . 000001 −p U200000, 200000 −hs L1 . 5, 1
−h l L1 . 2, 1 −hg l 1 . 4, 1 −u E10000000 −so U1, 1 −od 1 −or 0 −v 3
−cs 293745 −o model . 200 − 5 . 1000000 . 0 . 000001
~~~

For D2 with 1M generations we used

~~~
simphy −rs 50 − rl U1000, 1000 −rg 1 −s t U1000000, 1000000 − si U5, 5
− sl U200, 200 −sb U0 . 000001, 0 . 000001 −p U200000, 200000 −hs L1 . 5, 1
−h l L1 . 2, 1 −hg l 1 . 4, 1 −u E10000000 −so U1, 1 −od 1 −or 0 −v 3
−cs 293745 −o model . 200 − 5 . 1000000 . 0 . 000001
~~~

For D2 with 2M generations we used

~~~
simphy −rs 50 − rl U1000, 1 0 0 0 −rg 1 −s t U2000000, 2000000 − si U5, 5
− sl U200, 200 −sb U0 . 000001, 0 . 000001 −p U200000, 200000 −hs L1 . 5, 1
−h l L1 . 2, 1 −hg l 1 . 4, 1 −u E10000000 −so U1, 1 −od 1 −o r0 −v 3
−cs 293745 −o model . 200 − 5 . 2000000 . 0 . 000001
~~~

#### Gene length

*D1 (heterogeneous).* To draw the per-replicate *μ* and *σ* parameters of the log-normal distribution used for sequence length, we use the following approach. We first draw a number from a gamma distribution with shape of *k* = 2 log(1000) − 1/8 = 209.3259 and scale of *θ* = 0.033. Then we subtract this number from log(1000)/0.033 = 13.69051 to get the *μ*. The scale of the log normal distribution is also randomly drawn from a uniform distribution of (0.3, 0.7).
*D2 (homogeneous).* The log mean is drawn uniformly between 5.7 and 7.3, which correspond to 300 sites to 1500 sites. Thus, the average alignment length for each replicate is a random value between 300 and 1500. The log standard deviation for the log normal distribution is also drawn uniformly between 0.0 and 0.3. Base

